# Brain signatures of chronic gut inflammation

**DOI:** 10.1101/2022.10.22.513335

**Authors:** Caitlin V. Hall, Graham Radford-Smith, Emma Savage, Conor Robinson, Luca Cocchi, Rosalyn J. Moran

**Author notes:** **Correspondence to: Caitlin V. Hall**, QIMR Berghofer Medical Research Institute, 300 Herston Road, Brisbane, 4006, Australia.

## Abstract

Gut inflammation is thought to modify brain activity and behaviour via modulation of the gut-brain axis. However, how relapsing and remitting exposure to peripheral inflammation over the natural history of inflammatory bowel disease (IBD) contributes to altered brain dynamics is poorly understood. Here, we used electroencephalography (EEG) to characterise changes in spontaneous spatiotemporal brain states in Crohn’s Disease (CD) (n = 40) and Ulcerative Colitis (UC) (n = 30), compared to healthy individuals (n = 28). We first provide evidence of a significantly perturbed and heterogeneous microbial profile in CD, consistent with previous work showing enduring and long-standing dysbiosis in clinical remission. Results from our brain state assessment show that CD and UC exhibit alterations in the temporal properties of states implicating default-mode network, parietal, and visual regions, reflecting a shift in the predominance from externally to internally-oriented attentional modes. We investigated these dynamics at a finer sub-network resolution, showing a CD-specific and highly selective enhancement of connectivity between the insula and mPFC, regions implicated in cognitive-interoceptive appraisal mechanisms. Alongside overall higher anxiety scores in CD, we also provide preliminary support to suggest that the strength of chronic interoceptive hyper-signalling in the brain co-occurs with disease duration. Together, our results demonstrate that a long-standing diagnosis of CD is, in itself, a key factor in determining the risk of developing altered brain network signatures.

## INTRODUCTION

Immune dysfunction and accompanying systemic inflammation is thought to play a key role in the development of mood and affective symptoms (1, 2). As part of this mechanism, the presence of pro-inflammatory cytokines is communicated to the central nervous system (CNS) via peripheral activation of receptors expressed on vagal afferents, or the production of molecular intermediates at the blood-brain interface (c.f., circumventricular organs and the choroid plexus) (3). The brain recognises inflammation as a molecular signal of sickness, inducing changes at the neurophysiological and neurotransmitter level within brainstem, limbic and prefrontal regions (3, 4). Together, these neural responses generate a repertoire of “sickness behaviours” that includes social avoidance, anhedonia, fatigue, and depressed mood (1, 5, 6). The brain-cytokine response has been demonstrated in healthy adults administered lipopolysaccharides (LPS) (7) or typhoid vaccination (3, 8), who show transient alterations to cognitive-affective regions (involving the thalamus, amygdala, insula, and anterior cingulate), and a symptom profile that includes anxiety, poor mood, and impaired memory. These effects, however, embody the response of the resilient and adaptive CNS to an acute perturbation. Recent work investigating repeated exposure to immunogenic substances over an extended timeframe suggests more pervasive and enduring brain network abnormalities in chronic inflammation (9, 10).

Inflammatory bowel disease (IBD), a chronic, relapsing, and remitting intestinal disease, provides a unique and ecologically valid model to study the effects of inflammation chronicity on the brain (11). While IBD can occur at any age, disease incidence peaks in early adulthood (between 15 and 30 years) such that individuals experience a number of acute and recurrent inflammatory events that can endure for decades (12). As inflammation emerges within the gastrointestinal (GI) tract, the disease is well-placed to exert influence over the gut-brain axis (13). That is, the physical proximity of inflammation to the intestinal epithelium - a putative gut-brain interface - allows neural-related changes to be conceptualised as dysfunctions to vagal, immune, microbial, or endocrine signalling pathways. Alongside the mechanisms by which inflammation reaches and impacts the brain, an important research endeavour is focused on identifying specific brain regions affected by chronic inflammation, and how this can manifest behaviourally.

Neuroimaging work has provided initial insights into altered functional brain connectivity underpinning IBD pathophysiology, and suggests that a re-organisation of large-scale brain networks, rather than localised deficits, more clearly recapitulates disease-related changes (14–18). Specifically, there is a growing consensus that individuals with IBD exhibit alterations to default-mode network (DMN) activity (15, 18). The DMN comprises a set of brain regions that exhibit coherent neural activity during rest, and deactivation during externally oriented cognitive tasks (19). Alongside its involvement in social, cognitive and affective processes, the network plays a critical role in endogenous thought, such as rumination, and self-referential processing (20, 21). Abnormal patterns of activation and deactivation within the DMN has been linked to the development of neuropsychiatric disorders, including depression (22) and anxiety (23). In IBD, functional connectivity changes have been reported between key regions of the DMN, including the posterior cingulate cortex, medial prefrontal cortex, and precuneus regions (15, 18). Aberrant connectivity between nodes of the salience network (SN) (14, 17), including the anterior cingulate and insula cortex, further supports the possibility that IBD individuals experience altered interoceptive processing of visceral sensations (e.g., nociceptive, inflammatory, or microbial-related stimuli) (24). Given the relationship between the DMN and SN in anxiety and depression, the reported alterations in patients with IBD may be of substantial clinical importance. Critically, these results are reported in quiescent IBD (15, 16, 25–27), further supporting the argument that acute inflammation alone cannot account for the observed neural and behavioural impairments (9, 27, 28).

Among the two main IBD diagnoses, brain and behavioural abnormalities have more consistently been reported in Crohn’s Disease (CD) as opposed to Ulcerative Colitis (UC) (14, 15, 17, 18, 25, 26, 29). Despite overlapping symptoms, CD is thought to exhibit a more pervasive and severe disease expression attributed in part to the extent of affected anatomical sites, transmural involvement, and genetic and immune factors involved (30, 31). Moreover, while the microbiome in UC cannot be differentiated from controls following successful treatment, dysbiosis (imbalance) in CD persists long after remission and responds poorly to faecal microbiota transplantation (32–34). Despite well-defined heterogeneity between UC and CD - with the latter thought to express a more chronic and systemic disease profile – only a limited number of studies (35, 36) have directly compared IBD sub-groups in the context of whole-brain signatures.

In this study, we investigated whether CD and UC were associated with alterations to spontaneous brain state dynamics. To do this, we fit a Hidden Markov Model (HMM) to resting-state electroencephalography (EEG) data which describes brain dynamics as a sequence of transient and distinct patterns of power and phase-coupling within and between brain regions, respectively. We further explored these brain dynamics at a sub-network resolution, showing differential patterns of effective connectivity that are specific and selective to CD. Our results converge on the suggestion that long-term exposure to chronic gut inflammation confers a higher risk of altered brain and behavioural signatures, with the extent of these effects related to disease duration.

## METHODS

### Participants

The study was approved by the Human Research Ethics Committee of QIMR Berghofer Medical Research Institute (P3436). Written informed consent was obtained for all participants in accordance with the Declaration of Helsinki. Twenty-eight healthy controls (34 ± 11 years; 16 female), 40 CD (43 ± 13 years; 20 female), and 30 UC (42 ± 11 years; 21 female) participants were recruited from the Brisbane (Australia) metropolitan area by gastroenterologist (GRS) and accredited practising dietitian (CVH) (Supplementary Table 1). Exclusion criteria are presented in Supplementary Note 1. Study requirements involved (I) general health and clinical questionnaires; (II) neurocognitive assessments; (III) a resting state EEG recording; and (IV) a stool sample collected at home.

### General health and clinical questionnaires

The Brisbane Health Area Survey was administered to all participants and included questions about (I) current and previous medical history; (II) current and previous medical history of close family members; (III) medications taken in the previous 12 months; (IV) smoking, alcohol intake, and weight history; and (V) ancestry. The Traditional Mediterranean Diet (TMD) adherence questionnaire was administered by an APD (CVH). Prior to resting-state EEG recordings, blood pressure and heart rate were recorded. For individuals with CD and UC, an additional clinical questionnaire about IBD was administered, including detailed questions about (I) the nature and timing of symptoms experienced prior to a formal IBD diagnosis; (II) current and previous medications used to treat IBD; (III) current and previous history of procedures or surgeries performed in relation to their IBD; (IV) family history of IBD; and (V) comorbid health conditions associated with IBD, including extra-intestinal manifestations. Where available, the patient’s gastroenterologist provided clinical indicators of disease activity for CD (Harvey-Bradshaw Index, HBI) and UC (Simple Clinical Colitis Activity Index, SCCAI) patients, in a timeframe two weeks prior to, or two weeks post study participation.

### Neurocognitive assessments

Neurocognitive assessments were performed by a clinical psychologist and accredited practicing dietitian, and were used to rule out previous or current history of a neurological or psychiatric illness (excluding anxiety-related disorders or depression). Assessments of anxiety and depression included the Hamilton and Montgomery Anxiety (HAM-A), Montgomery-Åsberg Depression Rating Scale (MADRS), Hospital Anxiety and Depression Scale (HADS), Depression Anxiety and Stress Scale (42-item) (DASS), and Generalized Anxiety Disorder (7-item) (GAD-7).

### Sample collection and processing

Participants were provided with a stool nucleic acid collection and preservation tube (Norgen Biotek Corp., Thorold, Ontario, Canada) and were instructed to collect the sample within a window of 48 hours before/after the study session. Each stool sample was labelled and stored in a −80°C freezer until sample processing. Tissue homogenization was performed using tubes containing 1.4mm ceramic beads (Precellys Lysing Kit). DNA was extracted from samples and quantitated using Nanodrop 2000 (Thermo Scientific). PCR amplification was performed on the V3-V4 hypervariable region of the 16S rRNA gene, and sequenced on a MiSeq sequencer (Australian Genome Research Facility, Melbourne).

### 16S data processing and analysis

Demultiplexed fastq files were processed using default settings within QIIME2 2020.2 (https://qiime2.org) (37). Amplicon Sequence Variants (ASVs) were generated by denoising with DADA2 (38). For taxonomic structure analysis, taxonomy was assigned to ASVs using a pre-trained Naïve Bayes classifier and the q2-feature-classifier plugin against the Greengenes 13_8 99% 16S rRNA gene sequencing database. Samples were rarefied to a read depth of 2200 for diversity analyses. ANCOVA was used to test for group differences in Shannon diversity and Chao1 measures accounting for the effects of age, sex, and body mass index (BMI). Beta-diversity, assessed using unweighted UniFrac distance (39), was used to compare groups, controlling for age, sex, and BMI using qiime2 plugins PERMANOVA and adonis. The metagenomic functional contribution of each sample was predicted using the computational modelling approach, Phylogenetic Investigation of Communities by Reconstruction of Unobserved States 2.0 (PICRUSt2 v2.2.0-b) (40), using the MetaCyc Metabolic Pathway Database (41). The multivariate statistical framework, MaAsLin2 (42), implemented in R, was used to assess the relationship between group membership with (i) microbial abundance (collapsed at genus level) and (ii) functional pathway abundance. Covariates, including sex, age and BMI, were included as fixed effects. Features were included in if they had at least 10% non-zero values (across samples) and a minimum relative abundance threshold of 0.0001, both validated parameter settings in MaAsLin2. Significant features were corrected for multiple comparisons using the Benjamini-Hochberg FDR procedure, with corrected values of *p* < 0.05 and *q* < 0.25 considered statistically significant.

### Resting-state EEG recordings

Participants were fitted with a 64-channel EEG cap (Ant Neuro – EEGgo sports system), configured to the 10-20 international system. Signals were processed online using EEGgo with a sampling frequency of 2000 Hz. Scalp impedance was reduced to a maximum of 20 *k*Ω in all electrodes with the application of conductive gel. EEG activity was processed online using eego software. All electrodes were referenced to the CPz electrode. Prior to recordings, participants were reminded to keep their eyes open and fixate on a white crosshair against a black background. Participants were encouraged to breathe and blink normally, and relax head and neck muscles to minimize signal artifacts. Resting-state signals were recorded continuously for 4 minutes.

### EEG pre-processing

EEG data was pre-processed offline using EEGlab software (v2019.1) in MATLAB (vR2018b). The data were downsampled to 250 Hz. EEG signals were visually inspected, and excessively noisy channels were removed before signals were re-referenced to the common average reference (excluding EOG, M1 and M2 electrodes). Signals were band-pass filtered into a frequency band of 1-45 Hz, and epoched into 5-second segments. Epochs were manually inspected and removed if they contained large artefacts that would otherwise not be detected by independent components analysis (ICA) (e.g., strong muscle artifacts). Artifacts that were characteristic of cardiac, ocular or minor muscular movements were subsequently removed using ICA (InfoMax) (43). As the HMM is sensitive to noise, a fairly stringent approach was adopted to remove potential sources of signal artifact. This approach represents a necessary trade-off to ensure that the HMM is inferred on neurobiologically meaningful data and not spurious noise sources (44). As such, if more than 20 ICs were marked as artefactual, the original time series prior to ICA was re-inspected for additional sources of artefact. If more than 50% of epochs were removed, or more than 20 ICs were excluded after the second ICA run, recordings were excluded from the analysis. Recordings from 11 subjects (2 HC, 3 CD, and 6 UC) were not included in the HMM. Subsequent processing and analysis of EEG data were performed using toolboxes and software packages found within the Oxford Centre for Human Brain Activity (OHBA) Software Library (OSL) and SPM12. For source reconstruction, the forward model was generated using a symmetric boundary element method (BEM) and the inverse model was performed using a Linearly Constrained Minimum Variance (LCMV) vector beamformer. A 44-region weighted parcellation of the entire cortex was adapted from previous work (45–47). Thirty-eight parcels were constructed from an ICA of fMRI data from the Human Connectome Project, while the remaining six parcels corresponded to the anterior and posterior precuneus, bilateral intraparietal sulci, and bilateral insula cortex. The inclusion of the insula cortex - specific to our analyses - was based on previous work supporting the contribution of this region to interoceptive processing in chronic and inflammatory conditions, including IBD (15, 16, 48–50). Time-courses were extracted by taking the first principal component, with voxel contributions weighted by the parcellation. Symmetric multivariate spatial leakage (volume conduction) correction was applied (46).

### Time-delay embedded (TDE)-HMM

We adopted the TDE-HMM implemented within the HMM-MAR MATLAB toolbox (https://github.com/OHBA-analysis/HMM-MAR) (44, 45). We used stochastic variational Bayes (45) to infer the TDE-HMM parameterized with 6 states and 41 time lags (corresponding to a window length of 160ms) (Supplementary Fig. 1) using 500 training cycles and initialization parameters according to previously established procedures (45, 51–53). Prior to HMM inference, we concatenated time series across subjects from all three groups, producing a full dataset to obtain a common set of brain states across all participants. This approach facilitated a direct comparison of spatial and temporal statistics across groups (53, 54). Supplementary Note 2 provides a full description of the TDE-HMM and Supplementary Fig. 2 provides an overview of the analysis pipeline.

From the HMM we calculated the (subject-specific) temporal properties of each state using three parameters: (I) fractional occupancy, the proportion of total time spent in a state (K x 1); (II) interval time, the length of time between consecutive visits to the same state (K x 1); and (III) dwell time, the average length of time spent in a state before transitioning to another state (K x 1). We also computed subject-specific transition probability matrices representing the probabilities of transitioning from one state, to every other state (K x K). ANCOVA was used to test for significant differences in fractional occupancy, dwell times, and interval times between groups, controlling for the effects of age and sex. Permutation testing was used to reject the null hypothesis of equality between groups. As implemented in previous work (54), for each state we generated 5,000 permutations by shuffling group labels among participants. We then repeated ANCOVAs on the permuted values, therefore generating an empirical null distribution of *F*-statistics for each state and temporal measure (fractional occupancy, dwell times, and interval times). We ascribed statistical significance (*p* < 0.05) to the temporal values by assessing the proportion of null statistics that were greater than or equal to the value of the statistic computed for the non-permuted data. For significant ANCOVAs, Tukey’s HSD post-hoc paired t-tests were used to identify where differences were expressed between groups. The Network-based Statistic (NBS) (55) was used to perform inference on the transition probability matrices between the three groups, again including age and sex as covariates. We used an *F*-test with the primary statistic threshold set to 3.0, and performed a total of 5,000 permutations (family-wise error rate controlled at 5%).

### Candidate Cortical Regions

Using state-specific coherence values averaged across subjects, we calculated the eigenvector centrality (EC) measure for each region. EC calculates the centrality (degree) of each node and weights this according to the EC of the nodes that it connects to (56). EC was performed using the *eigenvector_centrality_und* function within the Brain Connectivity Toolbox (57). The top 10% of EC scores taken from a single hemisphere were used to inform regions for a DCM.

### Dynamic Causal Modelling

We used dynamic causal modelling (DCM) for cross-spectral densities (CSD) to selectively isolate those differences observed in the networks above (58, 59). Specifically, we modelled the extrinsic (between-region) effective connectivity strengths between candidate regions. We adopted the convolution based local field potential (LFP) neural mass model which describes source activity as the result of interactions between populations of inhibitory interneurons, excitatory spiny stellate cells, and excitatory pyramidal cells (60). The data to which the DCM was fit comprised the processed time series. For each subject, we specified and estimated a single model with a fully-connected network of 7 regions. To obtain the most robust estimates, we then re-estimated the DCM using an updated prior parameter space using the posteriors from an exemplar subject (Supplementary Fig. 3). For each subject, we selected the iteration with the best fit (as assessed by free energy). One-way MANCOVA (Wilks’ Lambda) was used to assess group differences in the forward and backward connectivity parameters. Univariate tests were corrected for multiple comparisons (*p_FWE_* < 0.05, Bonferroni corrected). A multiple regression model was used to assess the contributions of behavioural (non-clinical) variables to effective connectivity strengths.

## RESULTS

Resting-state EEG recordings and 16S rRNA profiles were analysed for 40 CD, 30 UC, and 28 healthy participants. Demographic, behavioural, and clinical characteristics are presented in Supplementary Table 1. IBD and healthy control (HC) participants were matched in terms of general demographics with the exception of age, and the Hamilton and Montgomery Anxiety (HAM-A) scores (Supplementary Table 1).

### Establishing distinct microbiota signatures in CD and UC

We first used 16S rRNA sequencing to compare microbiota profiles between the three groups. Our results show a significant difference in beta (unweighted UniFrac) and alpha (Shannon effective species and Chao1 index) diversity measures in CD, compared to HC and UC (**Fig. 1A-B**). While not reaching statistical significance, UC showed a trend towards lower alpha diversity and distinct beta diversity profiles compared to HC. Multivariate analyses also revealed a number of significant taxonomic and functional differences in CD and to a lesser extent, in UC (**Fig. 1C-D,** enlarged visualisation shown in Supplementary Fig. 4). The microbiota results converge in supporting the existence of a perturbed and heterogeneous microbial profile in CD (33). It is important to note that the small subset of CD participants exhibiting mild (*n* = 3) or moderate (*n* = 1, later excluded for poor quality EEG data) disease activity were not outliers in terms of their diversity scores (i.e., were distributed within the normal range for CD). Together, the clinical and microbiota results demonstrate clear distinctions between CD and UC sub-groups, providing a strong motivation to perform brain assessments in each group independently. Full statistical results for this assessment can be found in Supplementary Note 3.

**Figure 1.**
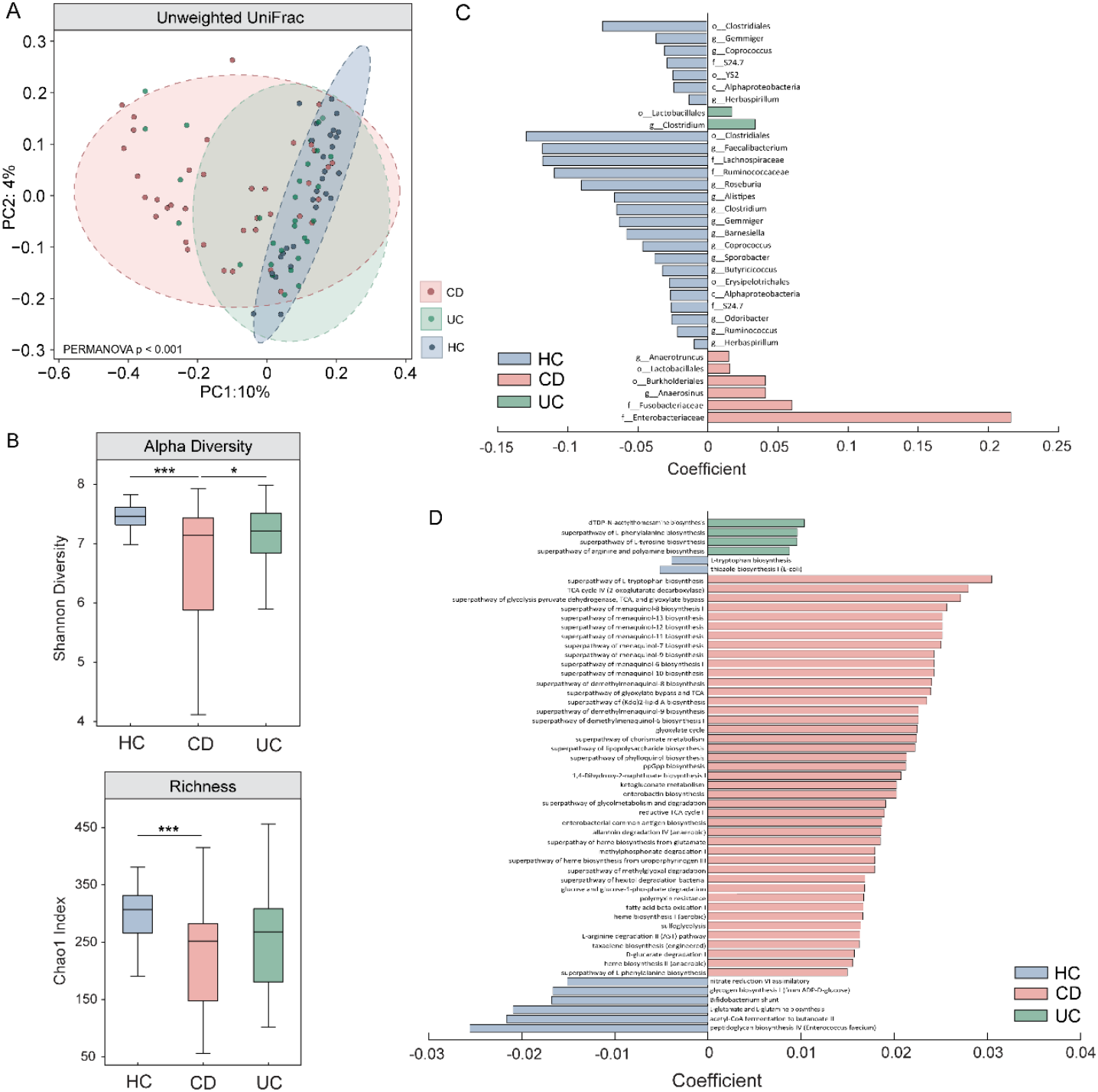
Comparison of microbiota results between Crohn’s Disease (CD), Ulcerative Colitis (UC), and healthy control individuals (HC). Results from (**A**) beta (unweighted Unifrac) and (**B**) alpha diversity (Shannon effective species and Chao1 index) measures show significant differences between CD and UC, and CD and HC, assessed using one-way ANCOVAs. Multivariate analyses performed using MaAslin2 revealed significant differences in (**C**) taxonomic abundance (genus resolution) and (**D**) functional pathways in CD and to a lesser extent, in UC, when compared to HC. Enlarged figures for (**C**) and (**D**) are presented in Supplementary Fig. 4. All microbiota assessments were controlled for the effects of age, sex, and BMI. * denotes *p* < 0.05; *** denotes *p* < 0.0005.

### Brain states expressed during resting-state EEG

We estimated brain states at rest using the TDE-HMM (61) (Supplementary Note 2). The HMM posits that a time series can be decomposed into a number of discrete and recurrent hidden brain states, comprising several regions that co-activate together, such that at each time point, only one state is active. Results showed that resting-state EEG data was best described by six short-lived and recurring brain states, each with unique spatial, spectral, and temporal profiles (**Fig. 2A-F**; Supplementary Fig. 5 and Supplementary Note 2). Our state selection is consistent with previous studies modelling M/EEG dynamics using the TDE-HMM, ranging between six and 16 states (44, 45, 62, 63). The spatial maps of power (i.e., the amount of activity) and coherence networks (i.e., the level of synchronisation or coupling between two regions) were averaged across a wideband frequency range (1-30 Hz). Power maps correspond to the mean power within each region and state (z-scored) and coherence networks show functional connections that are stronger (*p* < 0.01) compared to all other possible between-region connections for that state. Our spatial maps share characteristics with previous M/EEG HMM studies, including a bilateral pattern of activity for some, but not all states (44, 45), and strong increases in power often accompanying increases in coherence (45).

**Figure 2.**
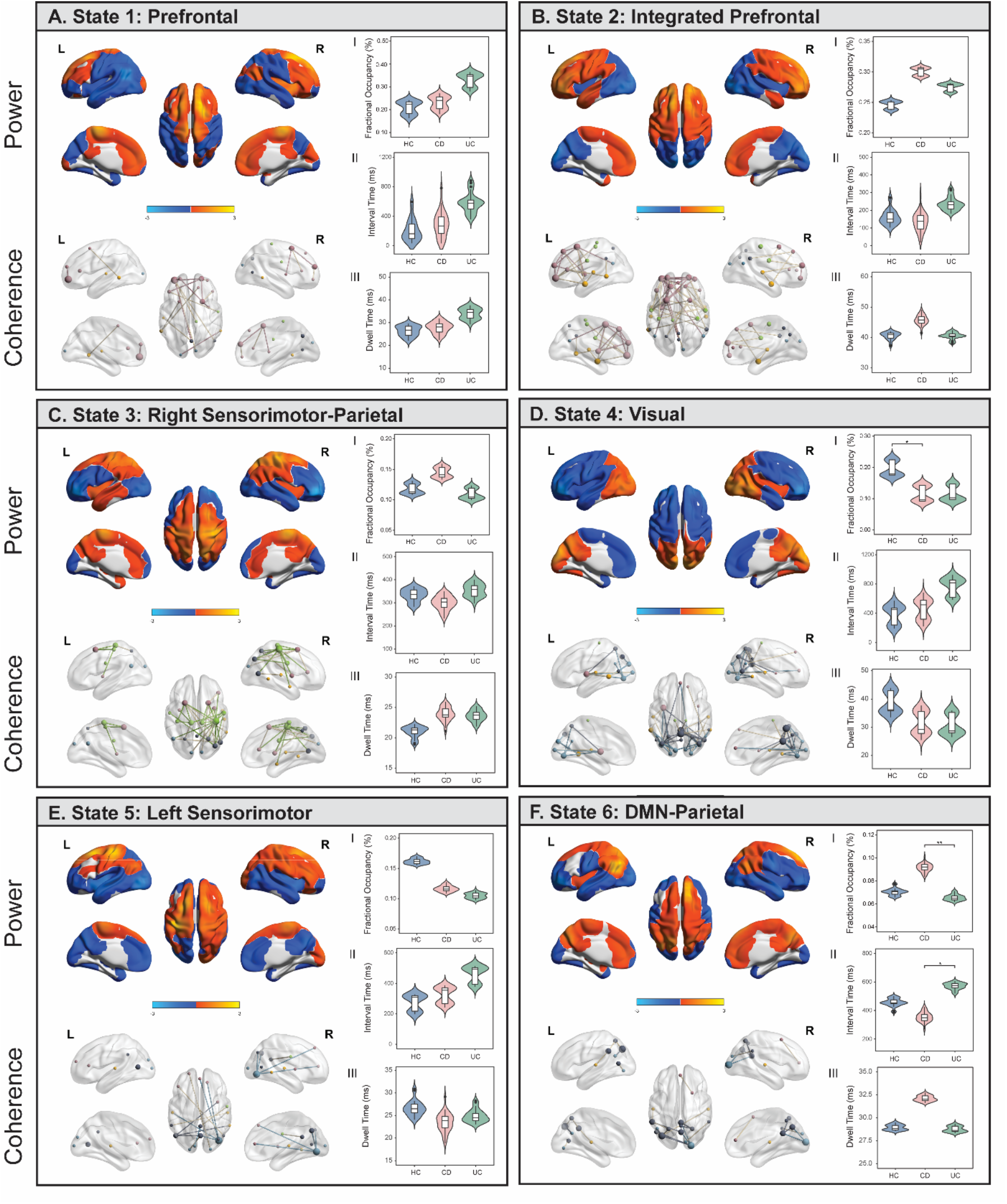
Brain states identified using Hidden Markov Modelling represent networks of power and spectral coherence. (**A**) Left panel shows wideband (1-30 Hz) power maps (top) and coherence networks (bottom) displayed for each state. Power maps are relative to the state average (z-scored) where blue colours reflect power that is lower than the state average and red/yellow colours reflect power that is higher than the average within that state. Coherence networks show statistically significant (*p* < 0.01) connections that stand out from a background level of connectivity within that state. Nodes are coloured based on which fMRI association map/s they anatomically correspond to, and the size of each node reflects the centrality (degree) score. (**A.I-III**) Comparison of temporal statistics between healthy controls (HC), Crohn’s Disease (CD) and Ulcerative Colitis (UC) individuals for each state, after adjusting for age and sex. Fractional occupancy (%) represents the proportion of overall time spent in a state; interval time (ms) represents the length of time between consecutive visits to the same state; and dwell time (ms) is the length of each state visit. Permutation tests were performed to assess the null hypothesis of equality in temporal measures between groups and Tukey’s HSD post-hoc tests were used to identify where significant pair-wise differences were expressed. * denotes *p_FWE_* < 0.05; ** denotes *p_FWE_* < 0.005.

### Brain states correspond to resting-state association maps

We quantified the functional overlap between the HMM states with established resting-state association networks from the meta-analysis database, Neurosynth (64). Specifically, we assessed the spatial overlap (voxel-wise correlation) between our power maps (z-scored, unthresholded) with canonical maps of prefrontal, parietal, sensorimotor, visual, DMN, and temporal fMRI association maps (Supplementary Fig. 6). For ease of interpretation, states were named according to the spatial patterns of activation to which they were most strongly correlated. States 1 (*prefrontal*) and 2 (*integrated prefrontal*) were defined by higher and lower power in prefrontal and visual regions respectively, with more extensive prefrontal coherence in State 2. States 3 (*right sensorimotor-parietal*) and 5 (*left sensorimotor*) were characterised by higher power in right and left sensorimotor regions, respectively, with coherence patterns closely following power in State 3. State 4 (*visual*) was characterised by high power and coherence in visual regions, while State 6 (*DMN-parietal*) reflected power and coherence in regions associated with DMN and parietal regions. Each state also exhibits frequency-specific differences in power and coherence, which can be visualized as an average across regions over the full spectrum (1-30 Hz) (Supplementary Fig. 5). There is a strong distinction between the *DMN-parietal*, characterised by power in the slower frequencies (delta/theta) and the *visual* state, characterised by stronger power in the alpha frequency. All states exhibit higher coherence within the alpha frequency band, with the strongest occurring in *right sensorimotor-parietal, visual*, and *DMN-parietal* states.

### Temporal brain state dynamics are differentially expressed in IBD

At each time point in the time series, the HMM estimates the probability that each brain state is active, referred to as the state time course. The state time courses estimated from the HMM were used to investigate between-group differences in three temporal statistics: (a) fractional occupancy, the proportion of overall time spent in a state; (b) dwell time, the length of each state visit; and (c) interval time, the length of time between consecutive visits to the same state. One-way ANCOVAs identified a significant main effect of group on fractional occupancy in the *visual* (*F*_(2,82)_ = 4.31, *p* = 0.018) and *DMN-parietal* (*F*_(2,82)_ = 7.40, *p* = 0.002) states, and a main effect of group on interval times in the *DMN-parietal* (*F*_(2,82)_ = 4.41, *p* = 0.001) (**Fig. 2D&F**). Relative to HC, individuals with CD resided for less time overall in the *visual* state (*p_FWE_* = 0.04) (**Fig. 2D-I**), but spent a longer time overall (*p_FWE_* = 0.002) and had shorter interval times between consecutive visits to the *DMN-parietal* state compared to UC (*p_FWE_* = 0.01) (**Fig. 2D-I-II**). While HC spent more time in the *visual* state compared to UC, and less time in the *DMN-parietal* state compared to CD, these effects did not survive Bonferroni correction (UC, *p_FWE_* = 0.08; CD, *p_FWE_* = 0.06).

We next used the probabilities associated with the state time courses to identify significant between-group differences in the transitions between brain states (Network-based Statistic (55), *P*_FWE_ < 0.05) (**Fig. 3**). Bold, coloured lines indicate transitions included in the significant NBS component while thin black lines show the top 20% most probable state transitions for each group (**Fig. 3B, D & F**). Firstly, individuals with UC were more likely to transition to the *prefrontal* state compared to HC and CD (**Fig. 3E-F**). Secondly, individuals with CD and UC were more likely to transition from the *left sensorimotor* to the *integrated prefrontal* state, while the inverse was true for HC. Finally, HC individuals were more likely to transition to the *visual* state, specifically from the *DMN-parietal* or *right sensorimotor-parietal* states.

**Figure 3.**
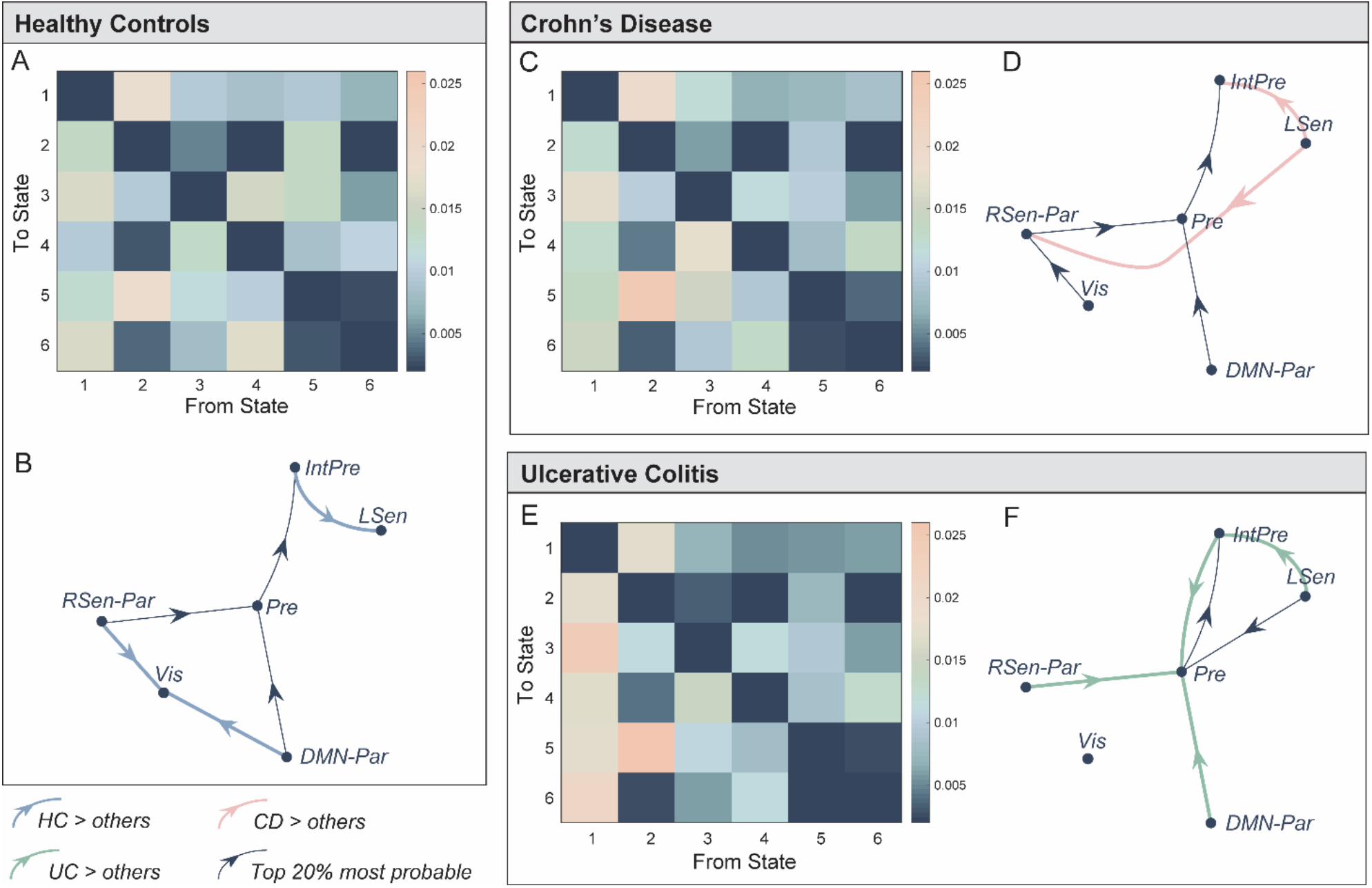
Representation of the transition probabilities between the six brain states in the three groups. (**A**) We computed subject-specific transition probability matrices representing the likelihood of transitioning from one state, to every other state (K x K). Diagonal matrix elements represent self-transitions (i.e., the probability of remaining in that state) and were set to zero to aid visualisation. (**B**) Directed transition diagram showing the top 20% most probable state transitions, where each arrow represents a transition. The thin black lines do not represent significant between-group differences, but represent transitions that were more probable on average for that group. Bold, coloured lines indicate a significantly higher probability of this transition in that group. The network-based statistics (NBS) was used to identify significant between-group differences in state transitions (*p_FWE_* < 0.05). (**C-F**) Same as (A) and (B) but for Crohn’s Disease and Ulcerative Colitis. *Prefrontal (Pre), integrated prefrontal (IntPre), right sensorimotor-parietal (RSen-Par), visual (Vis), left sensorimotor (LSen)*, and *DMN-parietal (DMN-Par)*.

Taken together, our results suggest that: (a) CD and UC individuals spent less time in, and are less likely to transition to the *visual* state; (b) individuals with UC are more likely to transition to, and may spend more time in the *prefrontal* state (although not reaching significance); and (c) individuals with CD reside for longer in, and spent less time between consecutive visits to the *DMN-parietal* state.

### Altered connectivity patterns between key brain regions differentiating groups

To identify the key drivers of these differences, we performed a refined sub-network analysis on communication between specific nodes within the *visual* and *DMN-parietal* states. We identified seven candidate regions exhibiting higher influence within each state’s spatial profile (See *Candidate Regions* in Materials and Methods). The posterior precuneus (Pprec), medial prefrontal cortex (mPFC) and left inferior parietal lobule (IPL) were identified within the *DMN-parietal* state, while the inferior occipital gyrus (IOG), mid occipital gyrus (MOG), and left insula (insula) were identified within the *visual* state (**Fig. 4A;** Supplementary Table 2). The posterior cingulate (PCC) had strong involvement within both states, supporting previous work recognising its “flexible” participation across a number of dynamic networks and associated cognitive processes (65, 66). Using the time series from each candidate region, we calculated the strength of directed (effective) connectivity, using dynamic causal modelling (DCM) (**Fig. 4B**) (See *DCM* in Methods and Supplementary Fig. 2 for details). Taking the expected values of the estimated connectivity parameters from all subjects, we identified a significant multivariate association between backward connectivity parameters and group membership (Wilks’ Lambda = 0.16, *F*_(84, 82)_ = 1.52, *p_FWE_* = 0.03). Univariate *F* tests identified a significant difference between groups in the connectivity from the left insula to mPFC (*F*_(2,84)_ = 8.57, *p_FWE_* = 0.017) (**Fig. 4C-D**). Specifically, individuals with CD showed significantly stronger connectivity from the left insula to mPFC compared to HC (*p* = 3.63 x 10^4^) and UC (*p* = 0.03). There were no significant differences between UC and HC for any connections.

**Figure 4.**
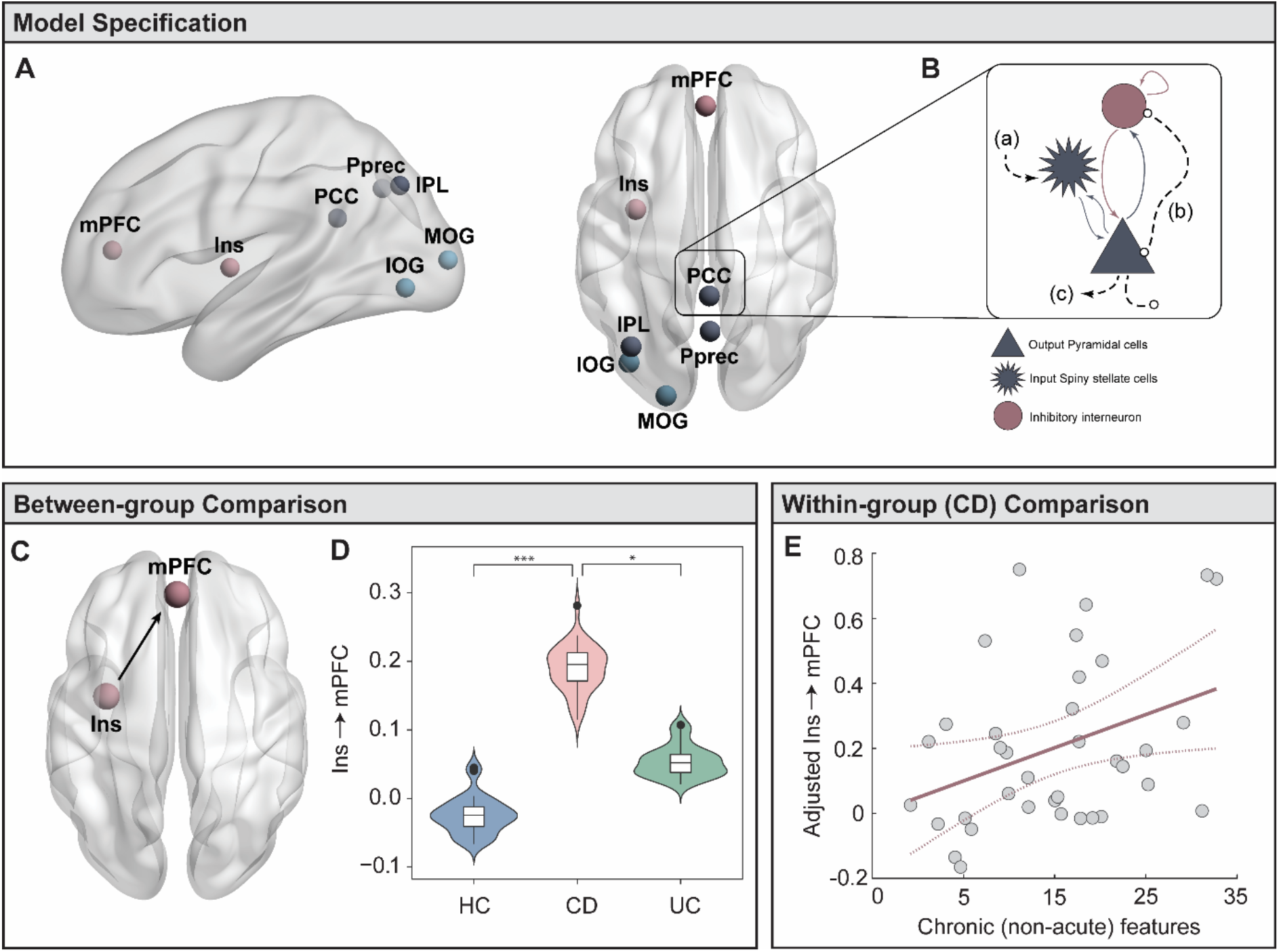
Targeted analyses of effective brain connectivity (Dynamic Causal Modelling, DCM). **(A)** Candidate regions were selected from the HMM brain states for a DCM analysis. (**B**) The Local Field Potential (LFP) convolution-based neural mass model was selected, modelling three subpopulations with five intrinsic connections. Extrinsic afferents are conceptualised as (a) forward connections arriving at the input spiny stellate population; (b) backward connections arriving at both the output pyramidal and interneuron populations. (c) Extrinsic (between-region) efferents project from the output pyramidal population to distant targets. (**C-D**) Results from MANCOVA post-hoc tests, showing significantly stronger effective connectivity from the left insula to the mPFC in CD individuals. (**E**) Multiple regression in CD group testing whether disease duration and behavioural symptoms predict the strength of left insula to mPFC connectivity. * denotes *p* < 0.05; *** denotes *p* < 0.0005.

### Insula to mPFC connectivity linked to disease duration in CD

Our results demonstrated a highly selective enhancement of connectivity between the left insula to mPFC in the CD group. With the exception of three individuals with mild disease activity, all CD individuals were in clinical remission. Thus, these findings provide support to our hypothesis that between-group connectivity differences may be driven by chronic, rather than acute inflammation. Our final aim was to specifically test whether inter-individual variability in the strength of insula to mPFC connectivity in CD was linked to long-standing disease features, thus testing for a more pronounced relationship of how CD chronicity (disease duration) links to depression, anxiety, and stress scores (DASS-42). The DASS-42 is based on a dimensional, rather than categorical, assessment of psychological symptoms, and provides higher inter-subject variability in sub-clinical populations. While the HAM-A and MADRS, and HADS-A and HADS-D, tend to produce anxiety and depression scores that are highly correlated, the DASS-42 is able to more clearly distinguish between anxiety and depression (67). Critical to our study, the anxiety scale assesses key components of interoceptive processing, including autonomic arousal, situational anxiety, and the subjective experience of anxious affect. While results did not show an overall multivariate relationship, we did find a significant independent regression coefficient linking longer CD duration (adjusted for participant age) with stronger insula to mPFC connectivity (*β* = 0.01, *t*_(34)_ = 2.19, *p* = 0.036) (**Fig. 4E**).

## DISCUSSION

In this study we assessed whether IBD - a model of chronic, relapsing, and remitting systemic inflammation - is associated with alterations in the spatiotemporal dynamics of spontaneous brain states. In particular, we directly compared CD and UC to delineate whether known distinctions in clinical, microbiome, and physical manifestations of gut inflammation also extends to variability in brain dynamics. Our findings extend upon previous work by showing a CD-specific brain signature implicating regions involved in cognitive-interoceptive appraisal mechanisms. The HMM assessment converges with these findings at a broader scale, demonstrating that IBD individuals exhibit alterations in the temporal properties of brain states supporting computations linking internal and external milieus. Together, our study supports a description of IBD as a dysfunction of the gut-brain axis, moving away from clinical definitions that compartmentalise effects in the gut from the CNS.

Our sub-network DCM dissected the relevant properties of the global brain state assessment. Specifically, this analysis provided a refined interpretation of global neuronal dynamics grounded in physiological and biophysical properties of the brain. These results showed a highly selective enhancement of connectivity from the insula to the mPFC in CD individuals (**Fig. 4C**). The insula is a key interoceptive hub, thought to be responsible for integrating information from the internal and external milieu to generate an awareness of the current emotional and internal state (68–70). During rest, information about the internal milieu likely emerges from gastrointestinal and cardiorespiratory stimuli before converging in the NTS and higher cortical regions, including the insula (24). Anatomically, the insula shares afferent and efferent connections with the mPFC (71) which together provide a contextual evaluation of emotional and affective states (72). The finding that CD individuals exhibit stronger bottom-up signalling from the insula to mPFC converges with a model describing altered interoceptive processing. As a function of persistent worry and rumination over anticipated visceral discomfort, many patients with GI disorders develop strong and rigid beliefs (i.e., hyperprecise priors) about the state of the body (73, 74). While the perception of abdominal pain in a healthy individual may not be considered alarming, the same signal may elicit hypervigilance in IBD. The perceived hypervigilance to visceral sensations has previously been cast within a predictive coding framework (75, 76). That is, the persistent inability to accurately detect afferent viscerosensory signals may produce a mismatch between top-down predicted states and the actual interoceptive input reaching the insula and prefrontal regions. This hypothesis is in line with a recent fMRI study showing altered interoceptive processing in CD to uncertainty about anticipated visceral discomfort, compared to controls (26). The tendency of an individual to overestimate the likelihood of a future aversive bodily state provides a conceptual bridge between altered interoception, and the development of clinical anxiety and depression (75, 77). While a confluence of factors are likely to contribute to the high prevalence of anxiety and depression in IBD, models describing the persistence and reinforcement of negative biases towards self-relevant information is thought to be a key contributor. As such, psychological interventions such as mindfulness and meditation have been put forth as adjuvant treatment approaches in IBD to modulate the brain’s response to future aversive interoceptive stimuli (78, 79).

Recent work has demonstrated that long-term exposure to recurrent systemic inflammation impacts brain and behavioural responses in a more permanent and pervasive way as opposed to a single inflammatory event (9, 10). Both CD and UC participants were either in clinical remission or had a mild disease course, with no difference in cardiovascular risk compared to healthy participants. Our results strongly suggest that brain dynamic alterations do not represent the effects of acute inflammation or vascular events, but suggests a more permanent network reconfiguration. In our study, we showed that insula to mPFC hyper-connectivity strengthens with disease duration in CD (**Fig. 4E**). The persistent and chronic effects from repeated exposure to inflammation are likely to result in a confluence of behavioural, biological, and neurophysiological changes, including alterations to interoceptive processing (e.g., heightened sensitivity to visceral inflammatory or nociceptive signals) (16, 26), hyper-activation of the hypothalamic-pituitary-adrenal axis (80), functional changes to the gut-brain interface, or altered serotonergic and glutamatergic neurotransmission (1, 81). In this study, we did not observe a relationship between effective connectivity and behavioural symptoms. However, it is possible that altered insula-mPFC hyper-connectivity represents a vulnerability towards developing psychological symptoms. Our results provide a strong motivation to pursue longitudinal assessments - monitoring fluctuations in inflammatory activity, medication use, symptoms, surgical procedures, and behaviour – to identify the causal mechanisms contributing to altered network signatures in long-standing CD.

Our results suggest that a diagnosis of CD is, in itself, a key factor in determining the risk of developing altered brain network signatures. Previous work suggests that UC and CD exhibit distinct disease processes (30, 31, 33). UC is described as a mucosal disease with an acute onset, while CD is considered a chronic and systemic disease with a long premorbid phase and transmural involvement (82). Systemic involvement in CD may also be reflected in the higher prevalence of extra-intestinal manifestations (83), with one study attributing low bone density to chronic and long-standing exposure to cytokines selectively in CD, but not in UC (82). Moreover, emerging work suggests that neurological effects related to IBD follows a differential pattern of involvement between sub-groups (84). That is, UC appears to exhibit extra-intestinal manifestations mostly in the peripheral nervous system, while CD is more closely associated with effects in the CNS (84). These observations are in line with a previous structural MRI (sMRI) and resting-state fMRI study comparing CD and UC sub-groups, showing that neural changes in CD may be more pronounced in patients exhibiting extra-intestinal manifestations (36). However in contrast to our results, this study, as well as another using near infrared spectroscopy (35), found that UC exhibited more pronounced neural changes overall compared to CD. Longitudinal and adequately powered studies will be critical to disentangle the nuanced alterations between CD and UC reported in this current study, and in previous work. Our results also showed that diversity, taxonomic, and functional microbiota profiles in CD are significantly different from UC and HC, despite the absence of major inflammatory activity (**Fig. 1**). The failure to restore eubiosis in CD may indirectly serve as a marker of chronicity, representing an epiphenomenon caused by repeated inflammation and extensive bowel damage, as well as a risk factor for recurrent relapse (32–34). Our observations underscore the broader significance of early diagnosis, and both rapid and effective control of gut and systemic inflammation in IBD patients.

A number of caveats need to be considered when interpreting the results from this study. The cross-sectional design and modest sample size are recognised as limitations. For example, our microbiota assessments of alpha and beta diversity did not detect significant differences between UC and HC. While these results are consistent with results from previous longitudinal 16S rRNA studies showing only diversity differences in CD compared to HC, but not in UC (33, 85), our relatively modest sample size may have resulted in a type II error (i.e., the non-detection of smaller effects in UC individuals). Secondly, our cross-sectional design does not allow us to disentangle the relative contribution of long-standing GI symptoms versus chronic inflammation to observed brain-related effects. However, recent works comparing UC to a control group with irritable bowel syndrome (GI symptoms without underlying inflammation) provides further support that these changes are more specifically driven by chronic gut inflammation, rather than long-term GI symptoms (50, 86). IBD is a heterogeneous disease, and even within CD and UC there is large variability in terms of surgical procedures, medication use, genetics, and inflammatory history. However, the main focus of this study was to characterise the large-scale brain effects from chronic, recurrent and relapsing gut inflammation within an ecologically valid and naturalistic setting. For example, a key source of heterogeneity was medication use in IBD. Specifically, there was a higher proportion of UC individuals taking aminosalcylates, analgesics, and corticosteroid medication (Supplementary Table 1). However, the fact that the CD group appear to be less reliant on medications overall – specifically analgesics – highlights that they have well-controlled symptoms. Taken together, this further strengthens the interpretation of our CD results, supporting the idea that hyper-connectivity from the insula to mPFC is more closely linked to disease chronicity, rather than acute inflammation or symptom flare-ups. Our study provides the initial impetus to pursue future targeted work, including a focus towards creating larger, longitudinal databases including multimodal neuroimaging, clinical, behavioural, and metagenomics data. While most IBD participants were in clinical remission, a number of participants were taking biologic agents, anti-inflammatory, or immunomodulatory medication. This suggests that some participants had experienced an acute inflammatory event at some point prior to the study. As we do not have longitudinal data about previous disease activity, we cannot directly assess their contribution to observed brain alterations. Unlike DCM, the TDE-HMM is a statistical method that is not grounded on biophysical models of neural activity. When inferring the HMM we recognize, like previous authors (45), that there is no biological ‘ground truth’ with regards to the number of brain states selected. Instead, varying the number of states simply offers different resolutions (spatiotemporal detail) to study brain dynamics. Selecting six states represented a necessary trade-off, allowing us to examine brain states that overlap with established fMRI resting state maps but in the process, limiting our ability to detect more subtle dynamics. Our HMM brain states were inferred from resting-state data. Future investigations could extend this work by assessing how external task-related demands modulate spatiotemporal dynamics in DMN and visual networks in CD and UC.

There is converging evidence showing the effects of acute inflammation on brain activity and behaviour (3, 7, 8). However, there remains a large gap in understanding how chronic and repeated exposure to systemic inflammation engenders change in spontaneous whole-brain dynamics. Using an ecologically valid model of peripheral inflammation, we demonstrate that CD individuals exhibit alterations in brain states and patterns of effective connectivity supporting computations within internal, interoceptive mental states. Our results provide motivation to pursue longitudinal assessments evaluating the impact of mood and affective disorders on the natural history of IBD, and vice versa. Understanding the extent and nature of gut-brain dysfunctions in IBD will help to optimise the monitoring and management of behavioural symptoms and critically, to prevent a gut disease from progressing to a comorbid psychiatric or neurodegenerative illness.

## DATA AVAILABILITY

Data supporting the findings of this study is available from the corresponding authors on reasonable request.

## ACKNOWLEDGEMENTS

We would like to thank L. Simms for their contributions to stool preparation.

## AUTHOR CONTRIBUTIONS

Conceptualization and methodology, Caitlin V. Hall, Rosalyn J. Moran and Luca Cocchi; Formal analysis: Caitlin V. Hall, and Rosalyn J. Moran; Data curation: Caitlin V. Hall, Conor Robinson, Emma Savage and Graham Radford-Smith; Writing—original draft, review, and editing, all authors; Funding acquisition, Graham Radford-Smith and Luca Cocchi; Supervision, Rosalyn J. Moran and Luca Cocchi.

