## Supplementary Material for "Brain signatures of chronic gut inflammation"

**Hall et al.**

### Supplementary Notes

#### Supplementary Note 1. Exclusion criteria for IBD and healthy controls

Exclusion criteria included: < 18 years old; a BMI of < 18.5 or > 30.0; current or previous history of a major psychiatric illness or neurological disorder (excluding depression and anxiety); major changes to dietary intake in the past month; consumption of ≥ 5 standard alcoholic drinks per day; recreational drug use ≥ 1 occasion in the past 3 months; acute disease (i.e., flu) at time of enrolment; pregnancy or lactation; chronic or clinically significant pulmonary, cardiovascular, hepatic, renal, or dermatological functional abnormality as determined by medical history; and history of cancer (excluding medically managed squamous or basal cell carcinomas of the skin). For healthy comparison subjects, additional exclusion criteria included history of an active, uncontrolled gastrointestinal condition, disease, or irregular bowel movements (including persistent diarrhoea or constipation); history of psoriasis or recurrent eczema; and use of the following medications within the past month: antibiotics, antifungals, antivirals, antiparasitics, corticosteroids, cytokines, methotrexate or immunosuppressive cytotoxic agents, or large doses of commercial probiotics.

#### Supplementary Note 2. Description of the TDE-HMM

We adopted the Time-delay Embedded HMM (TDE-HMM) implemented within the HMM-MAR MATLAB toolbox (https://github.com/OHBA-analysis/HMM-MAR), described in previous work[39,40]. Prior to HMM inference, we concatenated time series across subjects producing a full data-set to obtain a common set of brain states across all participants. This approach facilitates a direct comparison of spatial and temporal statistics across groups. As the source reconstruction via beamforming is performed individually for each subject, the signs of the reconstructed dipoles from the same parcel may be arbitrarily different across subjects. Thus, concatenating all subjects could result in suppressing or cancelling out phase relations between parcels. To mitigate these effects, a sign flipping algorithm[40] was applied in order to maximize the sign agreement across covariance matrices. The time series of each parcel was embedded with a time delay using *L* lags, resulting in an extended data matrix of (*L* lags * *N* parcels) * *S* time samples. Finally, to reduce computational demands and avoid overfitting, a principal components (PC) analysis was performed on the "embedded" space. We chose the number of PCs to be twice the number of sources (44 x 2), consistent with previous implementations[40,58].

**Selecting parameters for the TDE-HMM**

Two important parameters of the model that need to be specified a priori are the length of the window (i.e., number of lags) and the number of states. We used Stochastic Variational Bayes[40] to infer the TDE-HMM over a number of time lags (from $L$ = 11, to $L$ = 121, corresponding to 40-480ms) and states (from $K$ = 2, to $K$ = 14), using 500 training cycles and initialisation parameters according to previously established procedures[40,46-48]. Using the free energy as a formal comparison (and further re-represented as the percent change in Bayes Factor), we proceeded with the optimal model for these data - parameterized with $K$ = 6 and $L$ = 41 (with values between -20 and 20, corresponding to a window length of 160ms) (Supplementary Fig. 1). Using these parameters, the stochastic variational TDE-HMM inference was subsequently repeated 10 times, and downstream analysis proceeded on the model with the lowest free energy. We also performed an additional validation on state-specific temporal statistics, specifically, to ensure that (a) all states were represented in all subjects for at least a single time point; (b) no subjects were described by a single state; and (c) the mean fractional occupancy across subjects for any given state were not dominant (i.e., > 40% fractional occupancy).

**Temporal Properties of States**

To determine the temporal sequence of brain states, the HMM produces a state time course output that describes the probability of that state being active at each time point. The Viterbi path algorithm[77] then computes a non-probabilistic sequence of states (i.e., a hard-coding of the probabilistic state time course). The state time courses were used to calculate the temporal properties of each state between groups.

**Spatial Description of States**

The TDE-HMM is a variant that is inferred on raw time series (i.e., as opposed to power-envelopes[57]), allowing each brain state to be described by distinct patterns of power and phase-coupling. Wideband power and spectral coherence were extracted from each state using a previously validated state-wise multitaper approach[40]. To aid in the visualisation and interpretation of states, we factorised this information using the non-negative matrix factorisation (NNMF), so that power and coherence were averaged across all frequency bins, yielding coherence (44 x 44 x 6), and power (44 x 6) information for each region and state. The power maps were visualized according to each state's average (z-scored) such that blue colours reflect power that is lower than the state average, and red/yellow colours reflect power that is higher than the state average. For each state, we assessed which functional connections were significantly higher or lower relative to the other values within that state. To do this, we performed non-parametric permutations testing ($n$ = 5,000) on the spectral information calculated separately for each subject. Details regarding this assessment can be found elsewhere[40]. Coherence networks show the statistically significant ($p_{uncorr} < 0.01$) functional connections for each state. Using the wideband power maps, we also quantified the degree of functional overlap using association maps derived from the meta-analysis database Neurosynth[59]. This approach calculates the Pearson correlation between the two vectorized maps, where the r values reflect the correlation across all voxels between the two whole-brain maps.

**Contribution of higher frequencies to spatial descriptions**

To assess the contribution of higher frequencies (31-45 Hz) to the state-specific power and coherence differences, we performed the multitaper assessment on two wideband profiles: 1-45 Hz and a narrower frequency band (1-30 Hz). We observed that when using the narrower frequency band (1-30 Hz), regions with higher relative power tended to have higher nodal degree scores (i.e., there was greater overlap between power and coherence). This pattern has been identified in a previous TDE-HMM study[40], which they suggest may reflect the higher signal-to-noise ratio at these lower frequencies. Due to higher power-coherence consistency, and previous work suggesting large-scale patterns of synchronization at rest may be more relevant at lower frequencies, we used the narrower range to describe the spatial properties and downstream identification of candidate regions. However, we note candidate regions identified in the narrower frequency band were consistent with the 1-45 Hz band.

**Supplementary Note 3. Statistical details from the microbiota assessment**

We identified a significant difference in beta diversity (Unweighted Unifrac) between groups (*pseudo*-F = 2.22, $p$ = 0.001, *perms* = 999) (**Fig. 1A**). For alpha diversity, one-way ANCOVAs identified a significant main effect of group on diversity (Shannon: $F_{(2, 93)}$ = 9.28, $p$ = 0.0004) and richness (Chao1: $F_{(2, 93)}$ = 7.64, $p$ = 0.0013). Specifically, CD had significantly lower diversity and richness compared to HC (Shannon: $p = 2.74 \times 10^{-4}$; Chao1: $p = 8.39 \times 10^{-4}$), and lower diversity compared to UC (Shannon: $p$ = 0.02) (**Fig. 1B**). All analyses were adjusted for the effects of age, sex and BMI.

### Supplementary Figures

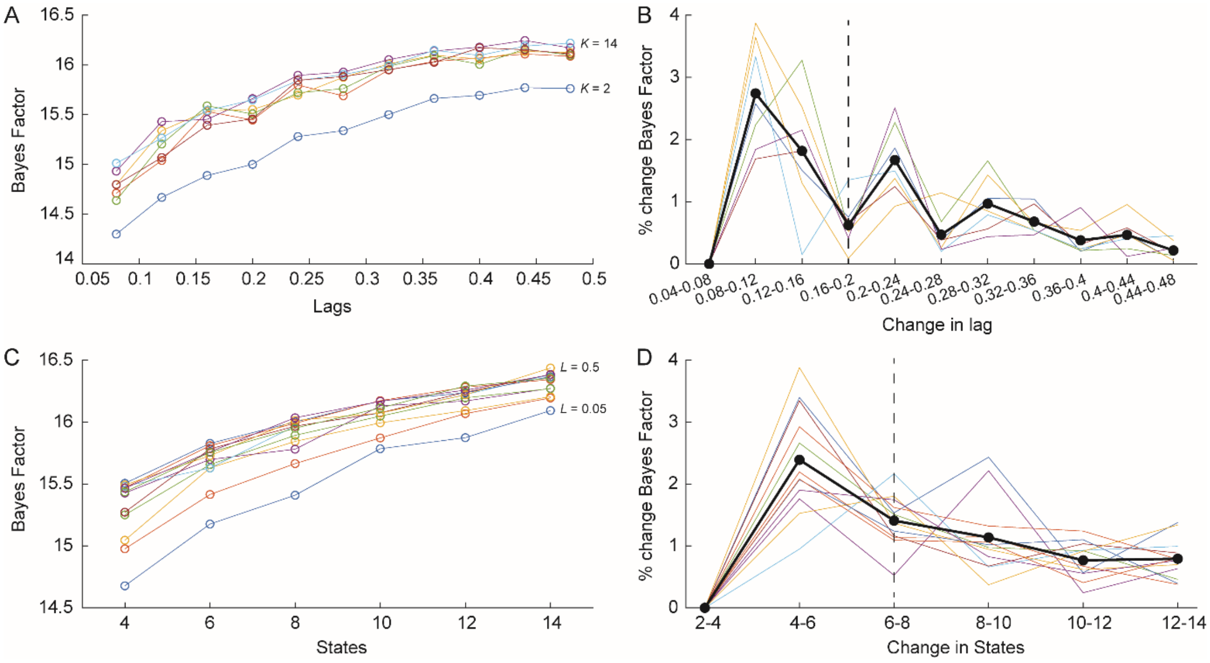


**Supplementary Fig. 1. Goodness of fit of the TDE-HMM.** The number of states (*K*) and the length of the lag (*L*) were parameters of the model that were specified a priori. Using stochastic variational Bayes, we inferred the TDE-HMM with a number of states varying from $K = 2$, to $K = 14$, and over a window length from $L = 40$ms, to $L = 480$ms, corresponding to 11 – 121 lags. (**A**) Variation in free energy (re-computed as Bayes Factor) and (**B**) percent change in Bayes Factor for each window length, where each line represents a different number of states. (**C**) Variation in free energy (re-computed as Bayes Factor) and (**B**) percent change in Bayes Factor across a number of states, where each line represents a different window length. Thick black lines represent mean percent change across states/lags, and dashed vertical black line indicates the selected state/lag used in final TDE-HMM analysis. The percent change decreases as the number of states increases beyond 6, and the window length extends beyond 160-200ms.

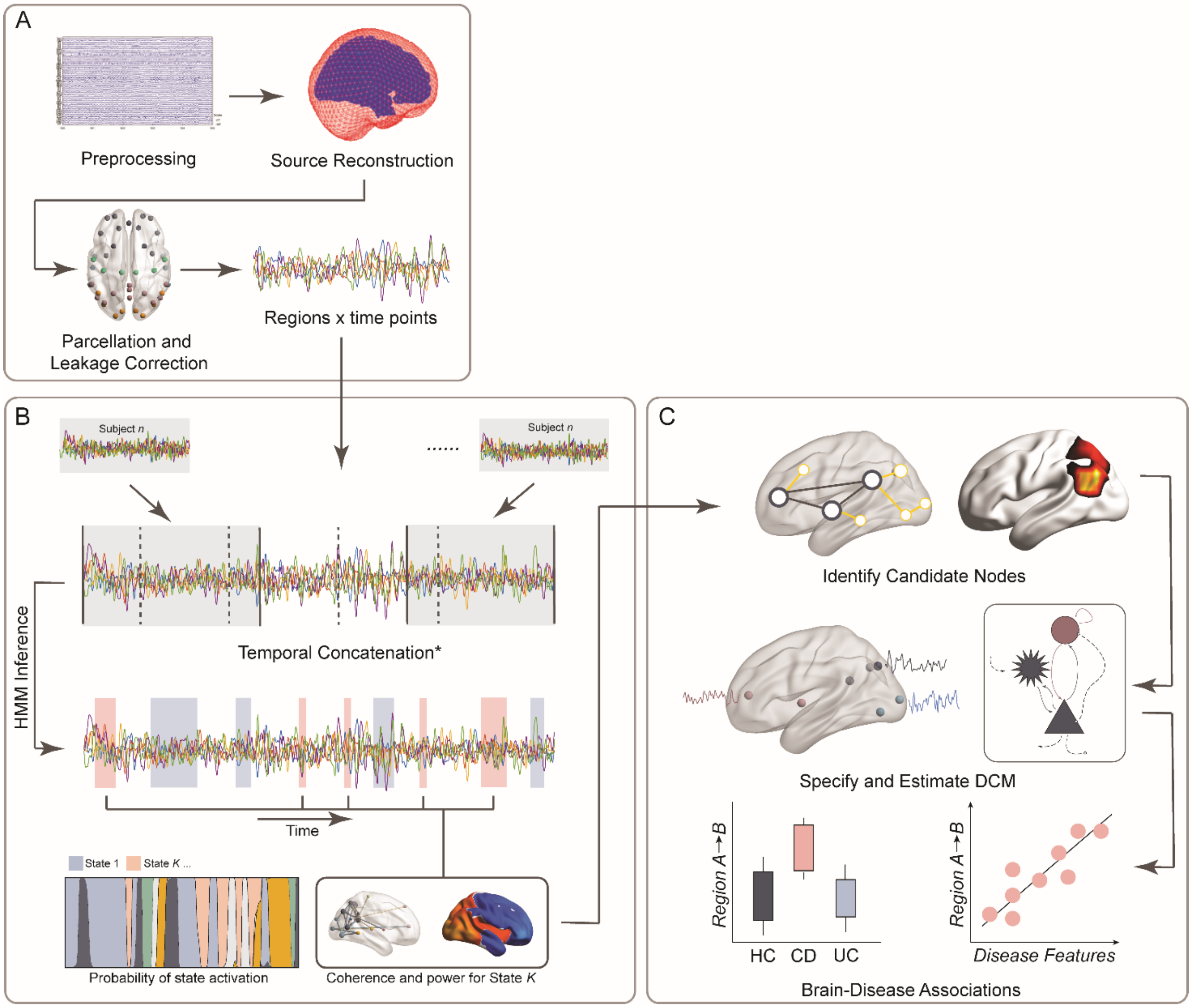


**Supplementary Fig. 2. Analysis Pipeline showing** (**A**) pre-processing of resting-state EEG data; (**B**) preparation and inference of the TDE-HMM; and (**C**) DCM analysis using candidate nodes informed from HMM brain state assessment. Between-group differences in effective connectivity were brought forward to an inter-individual assessment in CD only.

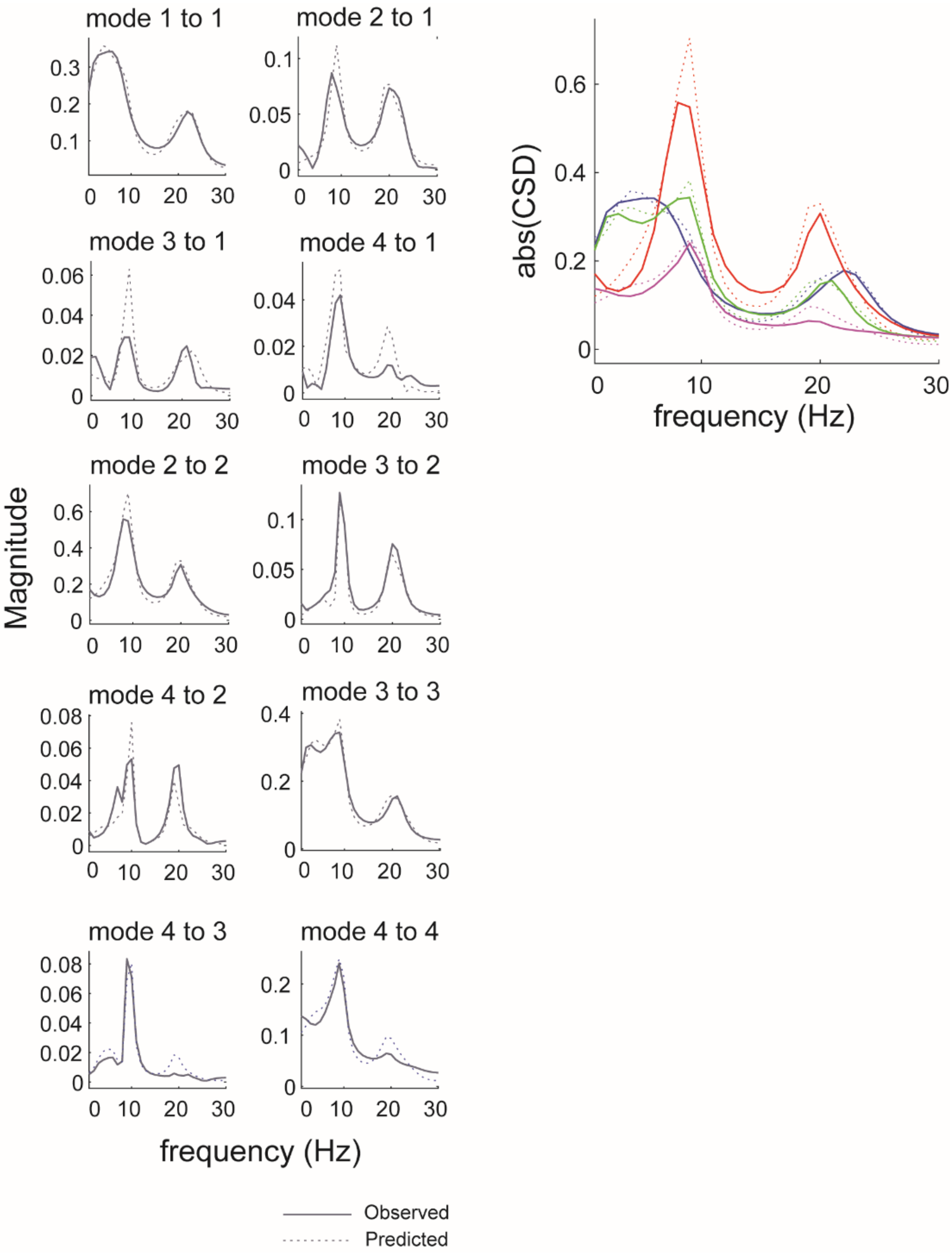


**Supplementary Fig. 3. Spectral Density over Modes.** Dynamic causal model (DCM)-predicted and observed auto- and cross spectra for the first 4 principal eigenmodes of EEG channel mixtures for a single exemplar subject. Mode 1 to 1, 2 to 2, etc. are auto-spectra while mode 1 to 2, 2 to 3, etc. are cross-spectra. The principal modes in our EEG data showed auto- and cross spectra often containing one or more marked peaks in alpha and beta bands.

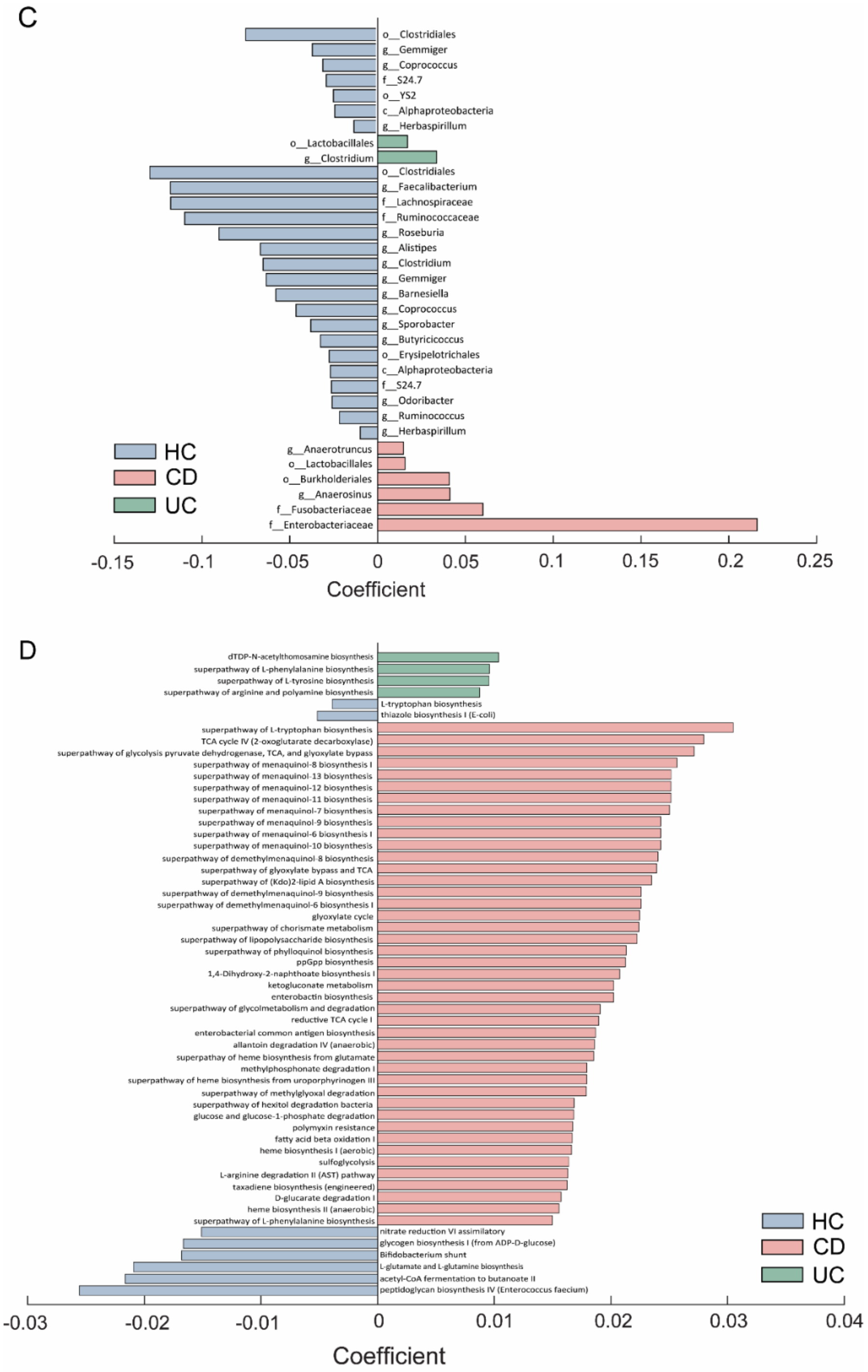


**Supplementary Fig. 4. Enlarged visualization of differential abundance testing between Crohn's Disease (CD), Ulcerative Colitis (UC), and healthy control individuals (HC).** Multivariate analyses performed using MaAslin2 revealed significant differences in (**C**) taxonomic abundance (genus resolution) and (**D**) functional pathways in CD and to a lesser extent, in UC, when compared to HC. Microbiota assessments were controlled for the effects of age, sex, and BMI.

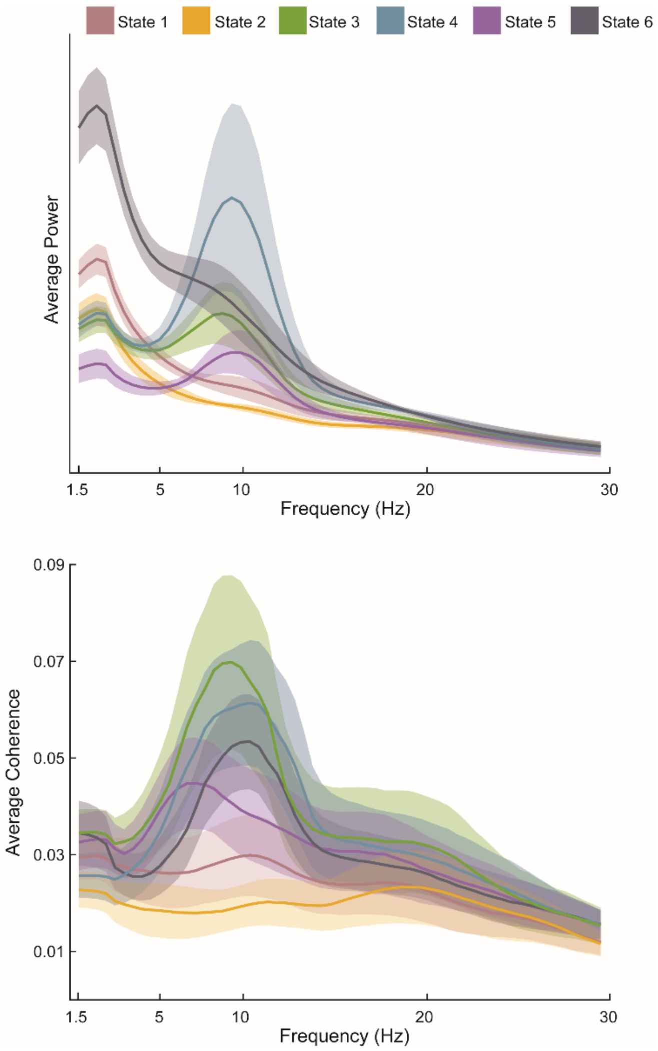


**Supplementary Fig. 5. Each state exhibits frequency-specific differences in power (top) and coherence (bottom), visualized as an average across regions over the full spectrum (1-30 Hz)**. There is a strong distinction between state 6 (*DMN-parietal*), characterised by power in the slower frequencies (delta/theta) and state 4 (*visual*), characterised by stronger power in the alpha frequency. All states exhibit higher coherence within the alpha frequency band, with the strongest occurring in state 3 (*right sensorimotor-parietal*), state 4 (*visual*), and state 6 (*DMN-parietal*) states.

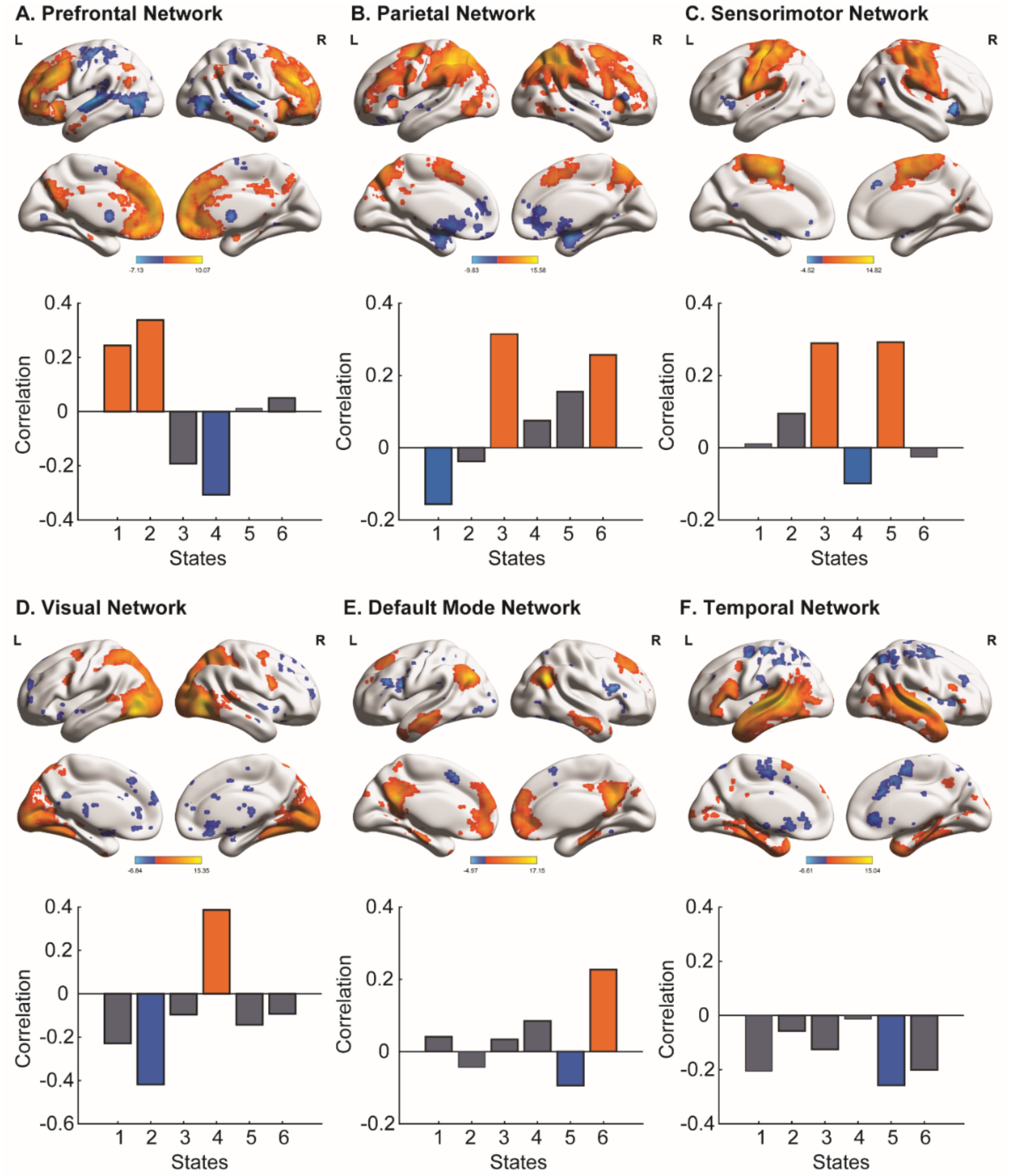


**Supplementary Fig. 6. Relationship between HMM brain states and fMRI association maps.** Using the meta-analysis database, Neurosynth[59], we calculated the Pearson correlation between the vectorized wideband power maps (unthresholded, z-scored) and the canonical fMRI association maps. That is, each HMM state is correlated with association maps representing (**A**) prefrontal, (**B**) parietal, (**C**) sensorimotor, (**D**) visual, (**E**) default-mode, and (**F**) temporal maps, where the r values represent the correlation across all voxels between the two whole-brain maps. The orange bars visualise the HMM brain state that more strongly correlates with the fMRI association map, whereas the blue bars show the HMM brain state that most negatively correlates with that fMRI association map. For ease of interpretation, HMM states were named according to the fMRI association map/s to which they were most strongly correlated.

### Supplementary Tables

#### Supplementary Table 1. Characteristics of study groups

| Demographics | CD | UC | HC | Between-group |
|---|---|---|---|---|
| Sample size (*N*) | 40 | 30 | 28 | - |
| Sex (male/female) | 20/20 | 21/9 | 12/16 | $X^2$ = 2.83, p = 0.24[a] |
| Age (years) | 43.2 ± 12.9 | 41.6 ± 11.3 | 34.0 ± 11 | **0.01**[b] |
| BMI (kg/$m^2$) | 24.9 ± 4.0 | 25.5 ± 3.9 | 24.0 ± 4.0 | 0.23[b] |
| SBP (mm Hg) | 123.3 ± 15.9 | 121.6 ± 14.4 | 121.3 ± 17.3 | 0.87[b] |
| DBP (mm Hg) | 81.5 ± 10.9 | 85.1 ± 7.7 | 83.5 ± 9.3 | 0.35[b] |
| Smokers (n) | 2 (NA = 8) | 0 (NA = 3) | 0 | - |
| Diet Score (TMD) | 6.2 ± 2.3 | 6.4 ± 2.1 | 6.9 ± 2.0 | 0.37[b] |
| Alcohol Intake (units/week, n) | Nil (4); ≤ 1 (17); 2-4 (3); 5-6 (3); 7-13 (7); 14-20 (2); 21-27 (0); N/A (4) | Nil (11); ≤ 1 (7); 2-4 (2); 5-6 (4); 7-13 (3); 14-20 (1); 21-27 (2); N/A (0) | Nil (1); ≤ 1 (5); 2-4 (5); 5-6 (6); 7-13 (5); 14-20 (0); 21-27 (1); N/A (5) | **0.05**[c] |

| Behavioural | CD | UC | HC | Between-group |
|---|---|---|---|---|
| HAM-A | 11.9 ± 8.1 | 9.0 ± 6.3 | 6.2 ± 6.5 | **0.01**[b] |
| MADRS | 9.9 ± 9.6 | 7.1 ± 6.1 | 5.5 ± 6.9 | 0.07[b] |
| HADS-A | 6.5 ± 4.4 | 6.0 ± 4.0 | 6.1 ± 4.0 | 0.88[b] |
| HADS-D | 4.2 ± 4.3 | 3.4 ± 2.9 | 3.0 ± 3.0 | 0.35[b] |
| DASS-D | 6.9 ± 9.3 | 3.7 ± 4.4 | 4.6 ± 7.8 | 0.21[b] |
| DASS-A | 5.0 ± 6.3 | 4.0 ± 4.7 | 4.3 ± 6.1 | 0.77[b] |
| DASS-S | 10.1 ± 8.0 | 10.5 ± 8.1 | 9.1 ± 7.1 | 0.79[b] |
| GAD-7 | 5.7 ±4.9 | 5.8 ± 5.0 | 5.1 ± 4.6 | 0.84[b] |

| Clinical | CD | UC | | Between-group |
|---|---|---|---|---|
| HBI, n | | | | |
| *Remission (< 5)* | 30 | - | | |
| *Mild (5-7)* | 3 | - | | |

| | CD | UC | Between-group |
|---|---|---|---|
| *Moderate (8 -16)* | 1 | - | |
| *NA* | 6 | - | |
| SCCAI, n | | | |
| *Remission (< 4)* | - | 23 | |
| *Active (5-19)* | - | 5 | |
| *NA* | - | 2 | |
| Disease duration (years) | 17.2 ± 11.1 | 13.5 ± 11.5 | 0.19[d] |
| Symptoms prior to dx (years) | 4.2 ± 6.3 | 2.8 ± 5.7 | 0.33[d] |
| Surgical History | | | 0.19[e] |
| *Pouch Procedure* | 1 | 4 | |
| *Colectomy Procedure* | 4 | 4 | 0.75[e] |
| *Ileostomy/Colostomy* | 8 | 1 | 0.08[e] |
| *Resection (Ileal, Jejunal, or Duodenal*) | 14 | 0 | **0.001**[e] |

| Current Medication | CD | UC | Between-group |
|---|---|---|---|
| Aminosalcylates | 2 | 14 | **0.002**[e] |
| Analgesics | 2 | 11 | **0.007**[e] |
| Antibiotics | 2 | 3 | 0.65[e] |
| Antihistamines | 2 | 5 | 0.23[e] |
| Biologics | 8 | 2 | 0.30[e] |
| Corticosteroids | 1 | 6 | **0.05**[e] |
| Immunosuppressants | 12 | 11 | 0.81[e] |
| Proton-pump inhibitors | 2 | 3 | 0.65[e] |
| No Medication | 15 | 6 | 0.31[e] |

*Mean ± SD; count (percentage); Median (range)*
[a] Pearson's Chi-squared test
[b] One-way ANOVA
[c] Kruskal-Wallis H test

[d] Two-tailed t-tests
[e] Fisher's exact test

**Abbreviations:** BMI, Body Mass Index; SBP, systolic blood pressure; DBP, diastolic blood pressure; TMD, Traditional Mediterranean Diet; HAM-A, Hamilton and Montgomery Anxiety; MADRS, Montgomery-Åsberg Depression Rating Scale; HADS, Hospital Anxiety and Depression Scale; DASS, Depression Anxiety and Stress Scale (42-item); GAD-7, Generalized Anxiety Disorder (7-item); HBI, Harvey-Bradshaw Index; SCCAI, Simple Clinical Colitis Activity Index; NA, data not available.

**Supplementary Table 2. MNI coordinates for candidate brain regions.**

| Region | Peak MNI Coordinates | | |
|---|---|---|---|
| | X | Y | Z |
| Posterior cingulate cortex | 1 | -49 | 24 |
| Posterior precuneus | 1 | -67 | 36 |
| Medial prefrontal cortex | 0 | 45 | 10 |
| Left inferior parietal lobule | -36 | -74 | 37 |
| Inferior occipital gyrus | -37 | -77 | -5 |
| Mid occipital gyrus | -20 | -94 | 7 |
| Left insula cortex | -34 | -4 | 3 |
